# Computationally Engineered CRISPR-SpyCas9 High-Fidelity Variants with Improved Specificity and Reduced Non-specific DNA Damage

**DOI:** 10.1101/2023.04.11.536265

**Authors:** Roy Rabinowitz, Oded Shor, Johanna Zerbib, Shay Herman, Natalie Zelikson, Shreyas Madiwale, Nataly Yom-Tov, Uri Ben-David, Felix Benninger, Daniel Offen

## Abstract

The CRISPR-Cas system holds great promise in the treatment of diseases caused by genetic variations. As wildtype SpyCas9 is known to generate many off-target effects, its use in the clinic remains controversial due to safety concerns. Several high-fidelity Cas9 variants with greater specificity have been developed using rational design and directed evolution. However, the enhancement of specificity by these methods is limited by factors like selection pressure and library diversity. Thus, *in-silico* protein engineering may provide a more efficient route for enhancing specificity, although computationally testing these proteins remains challenging. We recently demonstrated the advantage of normal mode analysis to simulate and predict the enzymatic function of SpyCas9 in the presence of mismatches. Here, we report several mathematical models describing the entropy and functionality relationships in the CRISPR-Cas9 system. We demonstrate the invariant characteristics of these models across different conformational structures. Based on these invariant models, we developed ComPE, a novel computational protein engineering method to modify the protein and measure the vibrational entropy of wildtype or variant SpyCas9 in complex with its sgRNA and target DNA. Using this platform, we discovered novel high-fidelity Cas9 variants with improved specificity. We functionally validated the improved specificity of four variants, and the intact on-target activity in one of them. Lastly, we demonstrate their reduced off-target editing and non-specific gRNA-independent DNA damage, highlighting their advantages for clinical applications. The described method could be applied to a wide range of proteins, from CRISPR-Cas orthologs to distinct proteins in any field where engineered proteins can improve biological processes.

## INTRODUCTION

The clustered regulatory interspaced short palindromic repeats (CRISPR)-Cas prokaryotic adaptive immune system^1^ was transformed into a powerful gene editing tool, allowing precise, efficient, and simple-to-design genome engineering^2,3^. Directed by a guide RNA (gRNA) molecule, the Cas effector protein binds the target sequence, subsequently leading to its cleavage or other editing alternatives^4–6^. The specificity of the CRISPR-Cas system has a clear impact on both genome-wide off-targeting and allele-specific targeting. Although being considered accurate, the CRISPR-Cas system has its limitations in mismatches discrimination^7–9^.

Previously reported engineered *Streptococcus pyogenes* Cas9 (SpyCas9) have been shown to have improved specificity. Each of these high-fidelity (HF) variants was engineered either by rational design^10–15^ or experimental directed evolution^16–21^. Rational design is limited to specific residues based on *a priori* knowledge. Furthermore, in most cases, it is constrained to alanine substitutions. Thus, the guiding principle is neutralizing residues and disruption of the protein activity in selected regions.

In contrast, directed evolution allows the freedom of random mutagenesis and evolvement of the protein by an iterative process of diversification and selection^22^. However, in addition to its time and labor-intensive nature, experimental directed evolution is commonly focused on specific regions of the protein or fails to cultivate potential candidates due to limitations of the host (i.e., bacterial or mammalian cells) and assay. Noteworthy, in many cases, substitution of residues distant from the active site of the protein, unexpectedly leads to significant changes in the protein’s activity^23^. Thus, constrained site-directed approaches will most probably overlook such impactful variants.

Comparable to processes that were transformed by digitalization, computational protein engineering could reduce time, labor and costs of research to become a powerful method. The challenge of transforming assays such as directed evolution or deep mutational scanning into a computational process, is the ability to assess the function of the evolving protein in the absence of living cells. Recently, researchers utilized machine learning to improve the on-target activity levels of KKH-*Staphylococcus aureus* Cas9 (SaCas9)^24^. Other methods were developed to improve the stability of proteins based on the energy differences between the reference protein and the variant^25,26^. Such tools were utilized to improve the activity levels of SpyCas9. However, together with the increased activity, the engineered variant demonstrated significant reduction in fidelity leading to elevated levels of off-targeting^27^. To date, no HF Cas variant has been engineered by a computational approach, plausibly due to the tradeoff between activity and specificity^28^. While the tools used to design the new variant aim to improve stability, it is likely that the enhanced stability of Cas9 will lead to compromised specificity. We hypothesized that the stability should be moderately reduced to improve the specificity of CRISPR-Cas proteins. Thus, to computationally design HF variants, it is necessary to employ a dynamic approach, capable of either increasing or decreasing the stability of the protein, in different circumstances and conditions, and simulating its functionality.

Recently, we demonstrated the ability to predict and simulate the activity of SpyCas9 in the presence of mismatches using normal mode analysis (NMA)^29^. The calculation basis of NMA involves a coarse-grained model that incorporates atom-specific interactions modulated by surface area contact and atom types, allowing for the prediction of protein dynamics and the effect of mutations on protein stability based on entropy considerations^30^. By calculating the vibrational entropy (*S_vib_*) of the SpyCas9 protein in complex with its single guide RNA (sgRNA) and target DNA, we obtained strong correlations between the NMA scores and the empirical SpyCas9 specificity profile^8^. Furthermore, we also demonstrated the compatibility of our approach for modified proteins, as we found a strong correlation between modelled known HF variants and their NMA scores^29^. In addition, NMA provides improvement in resolution compared to other methods to model structure-function relations, even when using coarse-grained NMA to overcome the computational limiting factor of analyzing large numbers of atoms in a protein^30–32^. Based on this success, we hypothesized that NMA could be utilized for protein engineering to obtain novel HF Cas9 variants by destabilization. In this work, we present a novel approach to protein engineering, termed Computational Protein Engineering (ComPE) mediated by NMA, which relies on *S_vib_* changes derived from a protein’s structure. Utilizing ComPE, we evolve SpyCas9 and pinpoint specific mutations that result in destabilization. We characterize nine candidate variants predicted by our ComPE method and experimentally validate their specificity attributes. Lastly, we demonstrate that HF Cas9 variants reduce gRNA-independent non-specific DNA damage, further enhancing the benefits of HF engineered variants in clinical applications.

## RESULTS

### NMA as the basis for the ComPE platform

NMA is a dynamic approach to study the function of proteins. We previously reported the use of NMA to link between genotype and phenotype in context of disease-causing mutations^33–36^, and its use to assess the specificity profile of SpyCas9 in the presence of mismatches^29^. Inspired by its ability to simulate the protein function, we used NMA to build ComPE, an *in-silico* method combining principles of directed evolution and deep mutational scanning. While experimental methods rely on survival or reporter-based assays, we first collected available data of engineered variants to establish the vibrational entropy (*S*_*vib*_) patterns in relation to experimental empirical data. The diversification stage has several parameters: fully or partially random, single or multiple substitutions, avoidance of active site mutagenesis, substitution to all or limited amino acids and other mutagenesis parameters. This stage goes from altering the primary structure of the protein to modeling the allosteric changes in the quaternary structure. Finally, we assessed the *S*_*vib*_ profile of each new generated structure using NMA and compared it to the generated *S*_*vib*_ profiles of known variants to identify potential improved candidates.

### Vibrational entropy of HF Cas9 variants correlate with the experimental empirical specificity

The association between the structure of a protein and its function is the underlying foundation of extrapolating the properties of the evolving protein. It is commonly accepted to classify Cas9 functions into on-target activity and specificity. While both parameters derive from the protein’s activity level, there is a clear negative correlation between the two. For example, HF variants with extremely high specificity (e.g., evoCas9), suffer from a severe reduction in on-target activity levels^20^. We used NMA to calculate the *S_vib_* of SpyCas9 and eight HF variants (eSpCas9(1.1), SpCas9-HF1, HiFi-Cas9, HypaCas9, evoCas9, xCas9, LZ3 Cas9 and Sniper-Cas9) and compared their whole protein mean *S_vib_* values to previously reported experimental measured specificity (**Fig 1A**)^20^. We noticed that the whole protein *S_vib_* of the WT and the eight variants related to the empirical specificity data could be described by the model function: 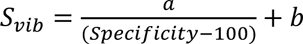. The goodness of the fit for the whole protein model function is *R*^2^ = 0.54. Considering that some individual residues show better fit with the model, we sought to assess the *S_vib_* of individual residues instead of the whole protein’s mean. For each residue in the protein, we measured the *S_vib_* and examined which residue returns the highest *R*^2^, representing the best fit for all nine Cas9 enzymes, between their *S_vib_* and specificity (**Fig. 1B**). It is noteworthy that the residue that showed the best fit to our model is not implied to be a candidate for mutagenesis, but rather a reliable indicator for the specificity of the enzyme based on the *S_vib_* (i.e., measuring the *S*_*vib*_ of residues with high *R*^2^ may provide a better indication for the function of the whole protein). We found that residue 632 had the best fit (*R*^2^ = 0.999). To formulate the function of the activity we had two assumptions; first, there is a negative relationship between activity and specificity. Furthermore, the empirical activity scores were normalized to WT. Thus, the following function represents the relationship between *S*_*vib* 1_ and activity: 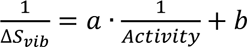. Here, _Δ*S*_*vib* is the normalization to WT: Δ*S*_*vib*_=*S*_*vib*_(*WT*) − *S*_*vib*_(*variant*). Due to the negative relationship between activity and specificity, their *S*_*vib*_ representations are reciprocal. Similar to the specificity analysis, we used the described function to explore individual residues and identify such that provide the best *R*^2^. Residue 350 provided the best fit with *R*^2^ = 0.94465 (**Fig. 1C**). To further improve the model, we tested various combinations of residues (**Fig. 1D-E**). The functions described above for specificity and activity were used to calculate the goodness of fit for each attribute, based on the mean *S*_*vib*_ of the residues combination. For the assessment of two residues, the *R*^2^ was calculated for the mean *S*_*vib*_ of every possible combination of two residues in the protein. Indeed, the goodness of fit for both models increased (*R*^2^[*Specificity*] = 0.99972, *R*^2^[*Activity*] = 0.988). We then tested every combination of three residues by taking the best hundred residues from the previous combinatorial analysis (combinations of two residues). We further took the best hundred residues from the last analysis and tested every combination of four residues. The same methodology was used to finally test every combination of five residues. This strategy resulted in *R*^2^∼0.99 for both models. These models, and the best combinations of five residues in particular, were used for our further analyses in this study.

**Figure 1:**
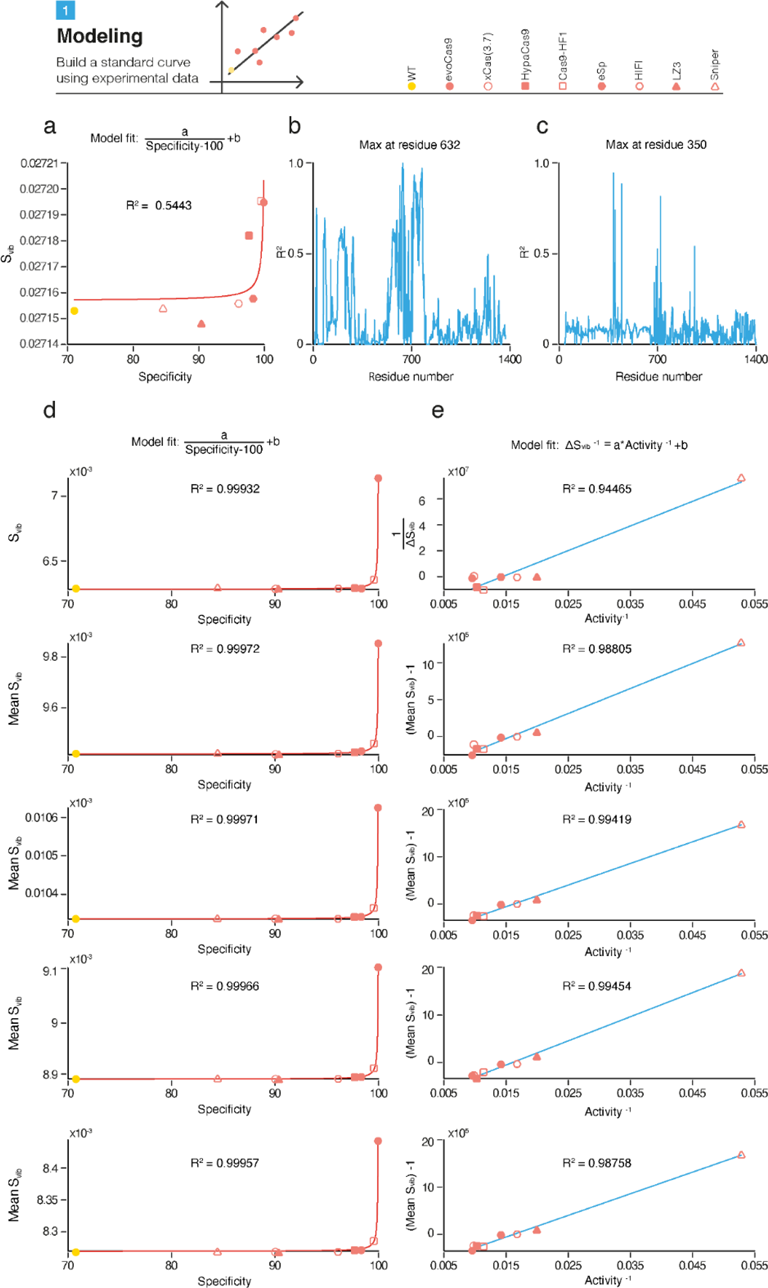
Identification of suitable models to depict the specificity and activity of SpyCas9 and HF variants using vibrational entropy. **a,** The interaction between the protein’s specificity and its vibrational entropy is described by the model: 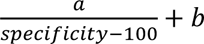, where the specificity values were obtained from Schmid-Burgk *et al.* ^20^ and the vibrational entropy was calculated using NMA. **b,** R^2^ represents the goodness of fit of the interaction between the specificity and *S*_*vib*_, calculated for each of the residues of all nine proteins to assess which single residue provides the best indication for all proteins. **c,** R^2^ represents the goodness of fit of the interaction between the activity and *S*_*vib*_, calculated for each of the residues of all nine proteins to assess which single residue provides the best indication for all proteins. The model fit for ^*S*^_*vib*_ is calculated as 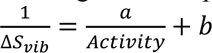. **d,** Improving the specificity-*S*_*vib*_ fit using combinations of residues. The mean *S*_*vib*_ represents the *S*_*vib*_mean of 2, 3, 4 or 5 residues, respectively. **e,** Improving the activity-*S*_*vib*_ fit using combinations of residues.

### The Robustness of the Model

To put the robustness of our approach to test, we sought to assess the effect of the starting empirical structural data. To that end, we selected SpyCas9 protein structures from various studies and conformational states (PDBs 4ZT0^37^, 6O0Y and 6O0Z^38^). The same methodology was applied on these starting models to test the goodness of fit. While the goodness of fit for the basic analysis of the best residue varied greatly, the lowest *R*^2^ obtained for the analysis of the best five residues was *R*^2^ = 0.955 (**Fig. 2A-B**). In addition, we used a series of protein structures demonstrating SpyCas9 bound to its on-target site and several off-targets^39^. Pacesa *et al.* reported the on- and off-target activity values in addition to the protein structures. These data allowed us to explore the interactions between the physical properties of the protein (i.e., entropy attributes) and the empirical protein’s function. Thus, in accordance with our previous study, we speculated that the relevant model should be:

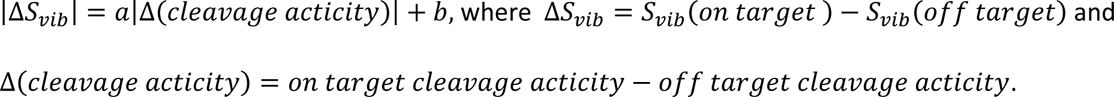

**Figure 2:**
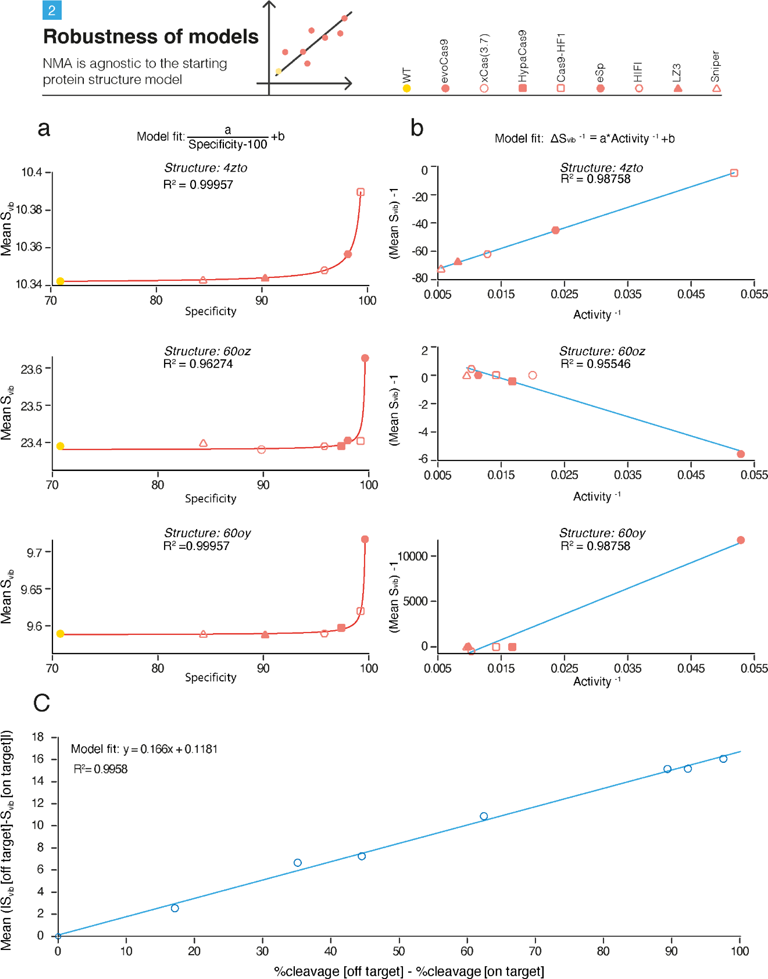
NMA is agnostic to the starting protein structure model. **a**, The modeling of the specificity-*S*_*vib*_ fit of WT and HF variants using three additional protein structures. **b,** The modeling of the activity-*S*_*vib*_ fit of HF variants using three additional protein structures. **c,** Correlation between empirical enzyme activity (obtained from Pacesa *et al.*^39^) and Δ*S*_*vib*_ = *S*_*vib*_(*on target*) − *S*_*vib*_(*off target*).

For a series of structures demonstrating SpyCas9 bound to its target sequence (FANCF, PDB: 7QQS) and seven additional off-target sites (PDBs: 7QQT, 7QQU, 7QQV, 7QQW, 7QQX, 7QR5 and 7QQZ^39^), we calculated the entropy of each residue in every structure using NMA. Then, by using the same combinatorial procedure described above, we obtained a three-residue combination that fitted well with our model and highly correlated to the reported on-target and off-target cleavage values. We observed a strong linear correlation and a very high fit with our model (*R*^2^ = 0.996) (**Fig. 2C**). These data provide strong evidence for the robustness of the method, indicating that our entropic-functionality models are independent on the conformational state of the protein depicted in the different empirical structural starting model. Furthermore, the comparison analysis of experimentally determined structures of SpyCas9 bound to off-target sites using NMA and empirical activity, emphasizes the ability of the method to simulate and predict the protein function in various genetic contexts. Taken together, these results suggest an invariant characteristic in the laws governing relations between entropy and functionality. Changes in the conformational structures do not alter the entropy-functionality mathematical model. The alteration of residues that fulfill the described mathematical relationship, across different conformational structures, resemble axis rotations and transformations observed in regular, conserved physical laws with symmetry. This analogy highlights the consistent relationship between entropy and functionality across various molecular structures.

### Diversification and Screening: In-silico deep mutational scanning and selection of HF Cas9 candidates

To discover novel single-mutation SpyCas9 variants with improved specificity, we diversified its structure by substituting each residue of the protein (1,362 residues) with every other amino acid (19 alternatives). In total, we generated and analyzed 25,878 new structures and preformed NMA to calculate their *S_vib_* profile, *S_vib_* for each residue in each of the structures. We defined the entropy score (ES) as the mean *S_vib_* values for the best combination of five residues that best fits our model. As anticipated, across all 25,878 single mutation structures, we found that the majority of single mutation substitutions resulted in minimal alterations to *ES*_*specificity*_, having most of the variants concentrated around *Median*_(*Activity*)_ = 8.21*x*10^−3^ (**Fig. 3A**) and *Median*_(*Specificity*)_ = 2.094*x*10^−2^ (**Fig. 3B**). Interestingly, previously reported HF variants have shown increased *ES*_*specificity*_ compared to WT SpyCas9, excluding Sniper-Cas9 showing lower *ES*_*specificity*_ on the specificity eCDF plot. Taken together, these results strengthen our hypothesis that moderate destabilization of Cas9, depicted with mild increase of *S_vib_*, would lead to improved specificity of the protein. We randomly selected nine candidates from the top 0.25% (*F*_(*entropy*)_ ≥ 0.9975), all characterized with higher *ES*_*specificity*_ than the known HF variants, predicted to have improved specificity (**Fig. 3C**), and mutated in different regions and domains of the protein. The selected candidates were not described in any previously reported HF variants: S512R, T519V, N692V, K735W, Q739R, V743N, K1129T, K1231I and R1359Q (in a sequence-chronological order). Noticeably, the *ES*_*specificity*_ of WT SpyCas9 was found to have *F*_(*Svib*)_ = ∼0.5 for both activity and specificity, meaning that about half of the single amino acid substitution variants have lower *ES*_*specificity*_ compared to WT SpyCas9, while the other half have higher *ES*_*specificity*_. The nature of the WT enzyme is to be stable and specific on one hand, but also adjustable to small changes on the other hand. Therefore, this observation may imply an evolutionary balance, leading to stability suitable for the function of the enzyme, and favorable over other small changes in the sequence of the protein. It is noteworthy that HypaCas9, which was generated by rational design, contains the N692A substitution along with three other alanine substitutions^12^. Our independent observation of the N692V candidate strengthens our hypothesis that increased *S_vib_* is the underlying mechanism of increased specificity in some HF Cas9 variants.

**Figure 3:**
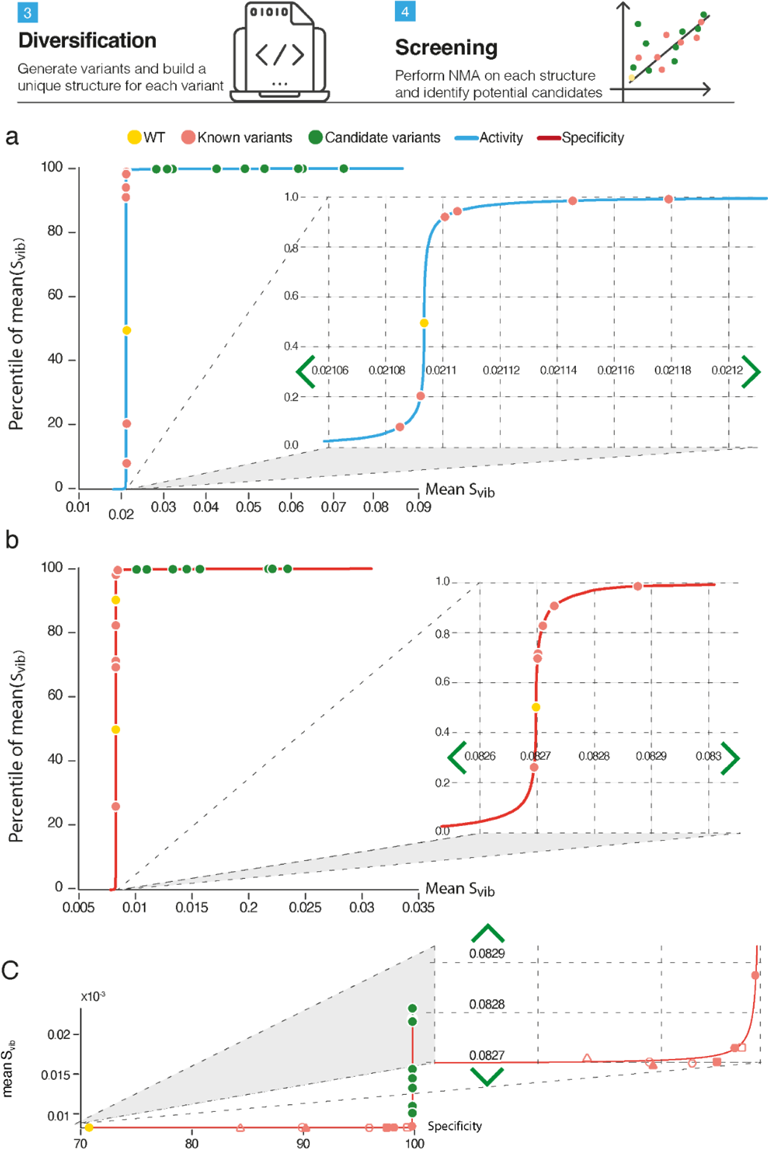
*In-silico* deep mutational scanning of SpyCas9 variants and analyses of their activity and specificity. **a,** eCDF plot of the proportion of variants and their log (|*S*_*vib*_|) values, based on the *S_vib_* score of the best five residues model fit analysis to assess the activity of all possible single-amino acid substitutions. The Y axis, *F*(*S*_*vib*_), describes the proportion of variants with a given value of log (|*S*_*vib*_|) or lower. The blue line represents the deep mutational scanning derivative variants (25,878 variants), while the dots represent WT (yellow) or the known HF Cas9 enzymes (light red) and the chosen candidates (green). The magnification box displays the sigmoid area of the curve. **b,** eCDF plot of the proportion of variants and their log (|*S*_*vib*_|) values, based on the *S_vib_* score of the best five residues model fit analysis to assess the specificity of all possible single-amino acid substitutions. The red line represents the deep mutational scanning derivative variants. **c,** Candidate variants with predicted improved specificity based on mean *S*_*vib*_.

### HF Cas9 candidates have reduced genome-wide off-targets

We then aimed to validate that the selected destabilized variants show increased specificity while conserving high activity of the enzyme. We generated a plasmid library of these nine candidates, as well as WT SpyCas9 and the previously reported HF variants eSpCas9(1.1), HiFi-Cas9 and Sniper-Cas9, into the same backbone, to eliminate differences due to different plasmid elements (i.e., regulatory elements and codon usage). We assessed the genome-wide off-targeting activity of our candidates compared to WT SpyCas9 and the known HF variants using a rhAmpSeq assay, a Next Generation Sequencing (NGS)-based method of on- and off-target evaluation using multiplex PCR^40^. We co-transfected any of the Cas plasmids with three sgRNAs targeting different loci with previously reported off-targets (EMX1, VEGFA and AAVS1)^20,40^ into HEK293FT cells, and extracted the genomic DNA 96 hours post-transfection. We first investigated the on-target activity of our candidate variants compared to WT SpyCas9, as many of the engineered HF variants are known to have reduced levels of activity^20,41^. Indeed, we observed decreased levels of activity in eSpCas9(1.1) and in some of our candidates (**Fig. 4a**). We then analyzed the off-target activity of the candidate variants, and identified that four of them (S512R, N692V, K735W and V743N) have significantly reduced off-target activity (**Fig. 4B, Sup. Fig. 1**). Moreover, we found reduced protein levels of the same candidates compared to WT and known HF variants, 72 hours following the transfection in HEK293FT cells (**Sup. Fig. 2**), demonstrating the lower stability of these variants. Together, these results highlight four novel SpyCas9 HF variants; S512R, N692V, K735W and V743N, and the N692V variant in particular, with both good on-target activity and reduced off-targeting. It is noteworthy that our data show that destabilization of SpyCas9 (**Sup. Fig. 2**) improves the specificity of the enzyme (**Fig. 4B**). However, we observed that previously reported HF variants (eSpCas9(1.1), HiFi-Cas9 and Sniper-Cas9) do not exhibit reduced stability compared to WT SpyCas9, although they demonstrated improved specificity. A plausible explanation would be the selection of Cas9 candidates with *S*_*vib*_ values higher than all other previously reported variants used to create the model. Thus, the increased *S*_*vib*_ of such variants might be moderate, enough to improve specificity, but insufficient to cause significant destabilization that can be detected on a western blot.

**Figure 4:**
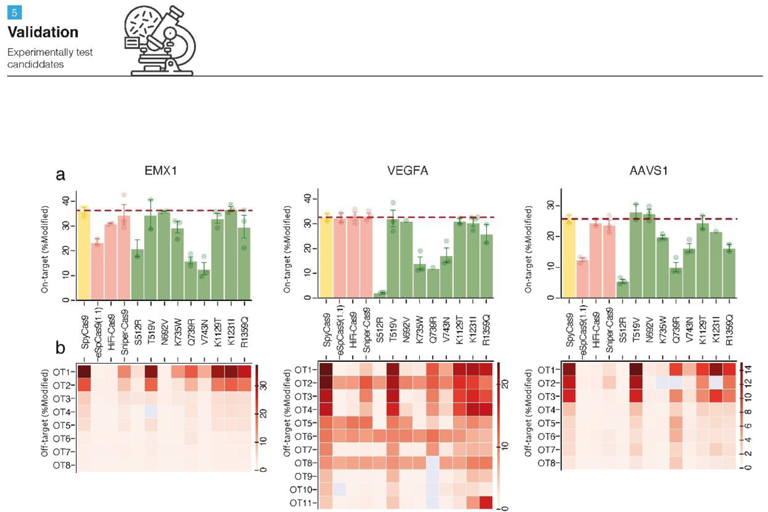
On-target activity and off-target analysis of the HF Cas9 variants. **a,** The on-target activity of WT SpyCas9 and the tested HF variants with three sgRNAs targeting EMX1, VEGFA and AAVS1. The activity is measured as the percentage of modified reads. The line indicates the average activity of WT SpyCas9. Error bars describe the standard error of the mean. **b,** Off-targeting by WT SpyCas9 and the tested HF variants with the three sgRNAs. Each off-target (OT) site is represented in a row. The white-red scale represents the percentage of modified reads for each locus. Only OT sites with >1% modified reads are shown. The data in both panels (a,b) represent three biological repeats for most samples. Grey boxes (b) are loci for which there were no reads (in any of the repeats).

### Single-base resolution – specificity profile of the HF Cas9 candidates

To further investigate our destabilized Cas9 candidates, we then characterized the specificity profile of each variant. In contrast to genome-wide off-targets analysis, the specificity profile describes the single-base resolution of Cas enzymes. Previous studies profiled the specificity profile of Cas enzymes using high-throughput methods, generally based on either flow cytometry^9,42,43^ or deep sequencing^8,44,45^. A high-throughput approach offers a deeper understanding of the mutational effects on the engineered variants, and it may explain the off-targeting patterns observed in unbiased methods such as rhAmpSeq, GUIDE-seq and others. Motivated by our previous off-targets analysis using rhAmpSeq, we employed a GFP disruption assay^9^ in EGFP-stable HEK293FT cells, using sgRNAs targeting the GFP gene with mismatches in each of the positions within the gRNA. We co-transfected the cells with both a Cas9 variant and an sgRNA and measured the activity of the Cas9 variant by quantifying the GFP fluorescence intensity using flow cytometry (**Fig. 5A**). We applied 20 different sgRNA combinations to each variant; we conserved one perfectly matched sgRNA to assess the maximal GFP reduction, while the other 19 sgRNAs carry a transversion mismatch (A↔T and G↔C) in positions 1-19. Since the hU6 promoter, which drives the expression of the sgRNA, requires a 5’G to initiate transcription, the 20^th^ position remained unaltered (**Sup. File 1**).

**Figure 5:**
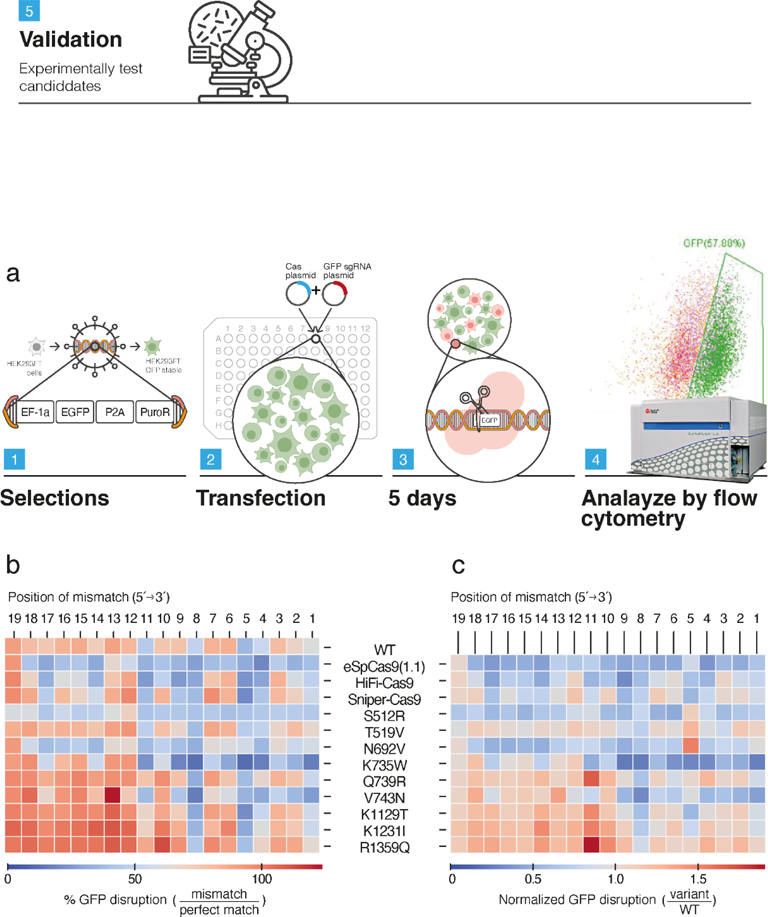
GFP-disruption assay reveals the specificity profile of HF SpyCas9 variants. **a,** Description of the experimental workflow. **b,** Heatmap representation of the GFP disruption by different Cas9 variants combined with mismatched gRNAs analyzed using flow cytometry. All values were calculated as the ratio of GFP disruption in the presence of a mismatch relative to the disruption with a perfect match gRNA. Thus, 100% indicates equal activity of the Cas9 variant combined with both the perfect match gRNA and in the presence of a mismatch. Values below 100% (towards the blue shades) indicate improved sensitivity to a mismatch in each position. Each data point represents the average of two biological replicates. **c,** Normalization of Figure 5c to demonstrate the sensitivity of Cas9 variants compared to WT SpyCas9. Values around 1 indicate no change in mismatch sensitivity compared to WT SpyCas9. Above 1 (dark red shades), sensitivity is impaired, while below 1 (blue shades), mismatch sensitivity is improved compared to WT SpyCas9.

Consistent with the rhAmpSeq results (**Fig. 4**), the four variants S512R, N692V, K735W and V743N exhibited improved sensitivity to mismatches, as demonstrated by the reduced EGFP disruption signals (**Fig. 5B**). We found different patterns of specificity between two groups of variants: the specificity of S512R and N692V was improved all along the gRNA, while K735W and V743N displayed improved mismatch discrimination along the seed region of the gRNA (positions one through eleven, **Sup. Fig. 3**). It is noteworthy that residues 512 and 692 are part of the REC III domain (REC lobe), while residues 735 and 743 lay within the RuvC II domain (NUC lobe) of *Spy*Cas9. Mutations of residues at both regions are commonly abundant in the variety of engineered Cas9 variants^7^. Furthermore, when we normalized the disruption created by each variant to WT Cas9, we found that S512R, N692V and K735W induced comparable disruption than the previously reported HF variants (**Fig. 5C, Sup. Fig. 3**). Overall, K735W demonstrated improved specificity in the rhAmpSeq assay and outperformed all other tested variants within the positions of the seed sequence in the GFP disruption assay. Taken together, these results validate the possibility to computationally obtain high specificity variants, better than WT SpyCas9, with comparable or even improved specificity compared to previously reported HF Cas9 variants.

### HF Cas9 variants reduce gRNA-independent activation of the p53 pathway

CRISPR-Cas9 was previously demonstrated to cause p53 activation in cell cultures^46–48^. Moreover, the introduction of Cas9 into cells, even in the absence of a gRNA, can result in non-specific DNA damage that leads to the activation of the p53 pathway^48^. We hypothesized that HF variants, and destabilized variants in particular, are less likely to undergo spontaneous conformational transitions, therefore less likely than WT SpyCas9 to cause non-specific DNA damage and activate the p53 pathway. To assess the effect of HF variants on p53 activation, we transfected p53-WT MCF7 cells with WT SpyCas9 or with HF variants and measured the expression of common p53-pathway downstream targets: GADD45, DDB2, RRM2B, XPC, TIGAR, MDM2 and CCGN1. As previously reported^48^, transfection of WT SpyCas9 significantly increased the expression of all tested p53 downstream target genes (**Fig. 6A**). We found that introduction of both previously reported (eSpCas9(1.1) and HiFi-Cas9) and destabilized (S512R and N692V) Cas9 variants significantly reduced p53 pathway activation compared to WT Cas9 (**Fig. 6A**), consistent with the reduced off-target and specificity profiles of all these variants (**Fig. 4**, **Fig. 5**). Summarized in a p53 transcriptional score, all tested variants exhibited a reduced activation of the p53-pathway compared to WT SpyCas9 (**Fig. 6B**). Immunoblotting of p21 also confirmed p53 pathway activation upon WT SpyCas9 introduction, while HiFi-Cas9 and our destabilized variants S512R and N692V demonstrated significant lower p21 activation, comparable to the stuffer plasmid control (**Fig. 6C-D**). Moreover, we found significantly less *γ*H2AX foci per nucleus in MCF7 cells transfected with HF variants than in cells transfected with WT SpyCas9 (**Fig. 6E-F, Supp. Fig 4A**), suggesting reduced accumulation of double strand breaks in HF transfected MCF7. Finally, p53 transcriptional score of each tested variant (**Fig. 6C**) significantly correlated (p=0.0292) with the number of *γ*H2AX foci per nucleus (**Supp. Fig. 4B**), confirming that HF variants induced less non-specific DNA damage than WT SpyCas9, emphasizing their importance in gene editing applications. Taken together, our results suggest that this destabilized variant may be of particular advantage for gene editing with minimal cellular damage.

**Figure 6:**
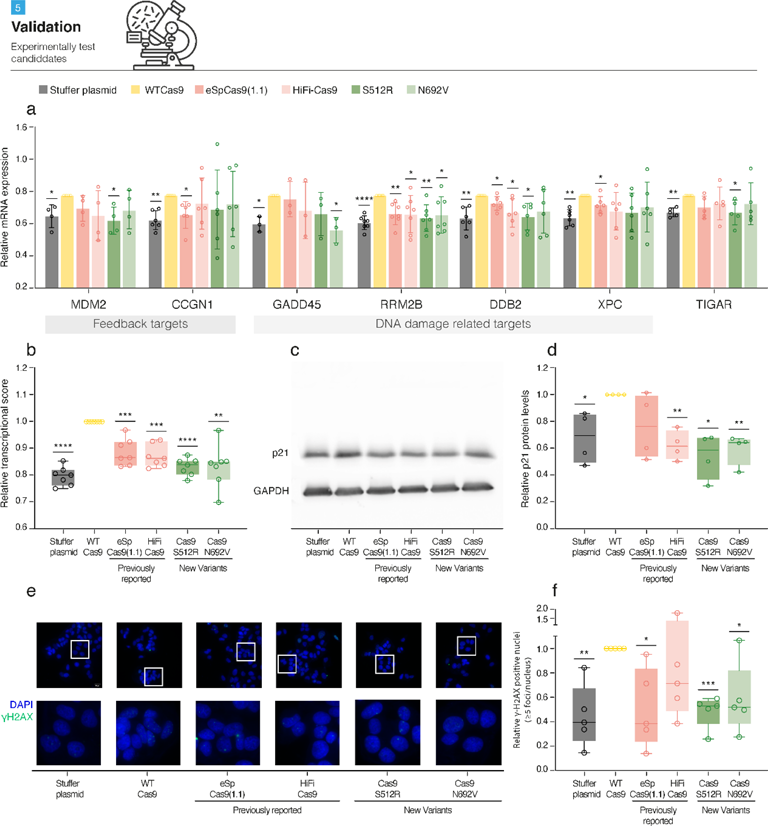
High fidelity variants activate the p53 pathway less than WT SpyCas9 and cause less non-specific DNA damage. **a,** Gene expression of multiple p53 downstream targets (MDM2, CCGN1, GADD45, RRM2B, DD2B, XPC and TIGAR) in MCF7 transfected with WT SpyCas9 or HF variants (eSpCas9(1.1), HiFi-Cas9, S512R and N692V). n=4 (MDM2), n=6 (CCGN1), n=3 (GADD45), n=7 (RRM2B), n=6 (DDB2, XPC), and n=5 (TIGAR) independent experiments. *p<0.05, **p<0.01, ****p<0.0001, One-Sample t-test. **b**, p53 transcriptional score for MCF7 transfected with WT SpyCas9 or HF variants (eSpCas9(1.1), HiFi-Cas9, S512R and N692V). Each dot represents the averaged gene expression for each p53 downstream target, as presented in Fig. 6a. n=7 targets, ***p=0.001 and p=0.0002 for eSpCas9(1.1) and HiFi-Cas9 respectively, ****p<0.0001 for Stuffer, S512R and N692V; One Sample t-test. **c**, Western Blot of p21 protein levels in MCF7 transfected with WT SpyCas9 or HF variants. GAPDH is used as a housekeeping gene. **d**, Western Blot quantification of p21 protein level, calculated relative to WT SpyCas9. N=4 independent experiments; *p=0.0438 and 0.0124 for Stuffer plasmid and S512R, respectively, **p=0.0063 and p=0.0061 for eSpCas9(1.1) and N692V respectively; One-Sample t-test. **e**, Immunofluorescence of *γ*H2AX foci in MCF7 transfected with WT SpyCas9 or HF variants. Scale bar 20μm. **f**, Quantification of *γ*H2AX foci in transfected MCF7 cells, calculated relative to WT SpyCas9. N=5 independent experiments; *p=0.027 and p=0.0335 for eSpCas9(1.1) and N692V respectively, **p=0.0085 and ***p=0.0009 for Stuffer plasmid and S512R, respectively; One-Sample t-test.

## DISCUSSION

We present ComPE, a novel approach for computational vibrational entropy-based deep mutational scanning. We employed ComPE to engineer SpyCas9 variants with improved specificity. While previously described Cas9 HF variants were engineered either by rational design or experimental directed evolution, here we report for the first time the use of an unbiased *in-silico* assay to predict and generate engineered variants with improved specificity (**Figs. 1-3**). Four (out of nine) of our predicted candidates were found to have improved specificity by experimental assays, pointing them as successful HF variants (**Figs. 4-5**). Notably, N692V also demonstrated both intact on-target and low off-target activities. As CRISPR-based therapeutics show promising results in clinical trials^49–52^ and move forward toward applicable solutions to treat genetic conditions, engineered Cas enzymes have a great potential to increase the safety of such treatments by reducing unintended off-target activity. Importantly, we also report that the use of HF variants leads to little-to-no non-specific DNA damage and p53-pathway activation (**Fig. 6**). The attenuated activation of the p53 pathway may reduce the selection for p53-inactivating mutations^48^, and also reduce undesired on-target effects, such as aneuploidy formation^53,54^. It may also have further benefits in homologous-directed repair (HDR)-based editing, as it is known that p53 inhibits the HDR process^46,47,55^. Previous studies also show that HF variants have increased HDR rates^56–58^. We propose that reduced activation of the p53 pathway may be responsible for HDR enhancement in HF variants editing. This hypothesis needs to be validated in a future study. Taken together, HF variants show great advantages for clinical and research applications.

Previous studies demonstrated that NMA could be utilized to analyze the effect of point mutations on proteins in terms of stability and flexibility^32,59^. Here, we report for the first time, the use of NMA to perform deep mutational scanning, based on reference experimental data, to improve a protein’s function in terms of binding specificity to binding counterparts (i.e., DNA, RNA, or other proteins). As binding properties can be described using entropy, we believe our approach is also relevant for engineering different biological macromolecules and their interactions. It is noteworthy that the mutations we report were non-trivial substitutions between different groups of amino acids with distinct properties (S512R, N692V, K735W and V743N), combining the best of both rational design and experimental directed evolution methods. Moreover, out of the four identified point mutations, two were found to be in close proximity to the gRNA or DNA (N692 and K735), while the other two (S512 and V743) are not in close contact with nucleic acids in the structure (**Sup. Fig. 5**). It is likely that allosteric effects account for the modified function of the protein. This observation emphasizes the impact of our method, similar to random directed evolution, but different from rational design: we identified beneficial mutations remote from the active site or ligand-binding residues. Another advantage of our computational mutagenesis approach is that the mutations are installed in the protein level (i.e., mutagenesis of amino acids in the structure), instead at the DNA level in experimental directed evolution. Our proposed amino-acid mutations (N↔V, T→V and K→W) required the alteration of two nucleotides in the codon, thus, are less likely to spontaneously arise during an experimental directed evolution campaign.

To affirm the reliability of ComPE, we examined a variety of protein structures featuring the SpyCas9 protein in different states (**Fig. 2A-B**). We also investigated structures of SpyCas9 bound to off-target sites, correlating the calculated vibrational entropy with the enzyme’s observed activity (**Fig. 2C**). The consistency observed across various starting models, states, and targets underlines the robustness of our method. Additionally, accumulated evidence from our prior research showcases the utility of NMA in examining diverse proteins and their functions through vibrational entropy^33–36^. We conclude that the identification of invariant relationship between the entropy and functionality upon different conformational structures strengthens the validity of the model and, in particular, the use of entropy for studying protein functionality. To the best of our knowledge, this is the first report proposing such an entropic invariant mathematical relationship in biological macromolecules. We propose asserting the validity of future models and methods in structural biology by examining invariance characteristics.

It can be assumed that in the future, advanced capabilities will enable multiple iterations of mutagenesis, enabling computational directed evolution and prediction of more complex combinatorial mutagenesis. Furthermore, availability of a protein structure is a limiting step for our method. This limitation might have been potentially solved by recent advances in protein structure prediction^60,61^. However, these and other tools are yet to be able to predict the structure of a complex with multiple counterparts (e.g., SpyCas9 bound to its sgRNA and target DNA), and their interactions. Such capabilities, combined with entropy-based methods, would greatly advance the field of computational protein engineering and drug discovery.

## METHODS

### NMA, entropy calculations and structure alterations

The structure of the SpyCas9 complex was taken from the Protein Data Bank (PDB-101; accession numbers PDB: 5F9R^62^). Using the mutagenesis plugin (default parameters) in PyMol Molecular Graphics System Version 1.8 (Schrödinger, LLC., Cambridge, MA), we created all possible variant structures with single amino acid residue mutagenesis (total of 25,878 single residue mutation structures), as well as the eight previously reported HF variants. The WT and each of the possible single residue mutated structures were analyzed using ENCoM coarse-grained NMA to evaluate the effect of the mutation on the vibrational entropy of the protein. This method is based on the entropic considerations C package of ENCoM^31^ available at the ENCoM development website (https://github.com/NRGlab/ENCoM), compiled and used on an Ubuntu platform (Canonical Group, UK). We employed ENCoM for the NMA due to the report that ENCoM outperformed other entropy-based methods to predict the stabilizing effect of mutations^30^. For each analyzed variant, we calculated the vibrational entropy profile of the structure. The same approach was taken to generate the known HF variants to establish the correlation curves. Finding the most correlative residue vibrational entropy of the six high fidelity structures and their activity or specificity scores was performed using MATLAB software (MathWorks, Natick, MA).

### Cell culture, transduction and transfection

HEK293FT cells were grown in DMEM supplemented with 2% L-glu, (Gibco 41965039), 10% Fetal Bovine Serum (Gibco 10270-106) and 1% Penicillin-Streptomycin (Gibco 15070063). MCF7 cells were grown in RPMI (Sartorius) supplemented with 4mM Glutamine (Sartorius), 10% Fetal Bovine Serum (Gibco) and 1% Penicillin-Streptomycin (Sigma). All cells were maintained at 37 °C with 5% CO2.

To establish the EGFP-PEST stable cell line, HEK293FT cells were transduced with a lentivirus carrying the EGFP-PEST-P2A-PuroR construct. Following transduction, infected cells were selected with puromycin (Thermo-Fisher J67236, 1:1000) for two weeks. For Cas9 activity assays (rhAmpSeq and GFP disruption), HEK293FT cells were transfected with 40ng plasmid DNA/well (sgRNA:Cas ratio 1:1, supplemented with mCherry reporter plasmid for transfection assessment and flow cytometry gating), using 0.2ul/well Lipofectamine3000 (Invitrogen L3000015) and OptiMEM (Gibco 31985-047), following manufacturer’s protocol. All transfections were carried in 96-well plates using 8,000 cells/well. Transfection media were replaced 24hr post-transfection.

EMX1 and VEGFA site 3 gRNA sequences were cloned into BPK1520^63^ backbone according the standard protocol. All other gRNAs and Cas enzymes were cloned into expression vectors by VectorBuilder.

Estimation of p53 pathway activation following introduction of Cas9 variants was performed as previously described^48^. Briefly, MCF7 cells were transfected with 2ug of DNA constructs using Trans*IT*-LT1 reagent (Mirus), following manufacturer’s protocol. RNA was extracted 72hrs post transfection.

### DNA extraction

For rhAmpSeq analyses DNA was extracted 96hrs post transfection, as previously described^64^. Briefly, cells were carefully washed with PBS and then lysed with 150ul lysis buffer (10mM Tris-HCl pH8, Sigma T3038; 0.05% SDS, Bio-Basic SD8119; 35ug/ml Proteinase K, Roche 03-115-879-001). After 1hr incubation at 37 °C, lysed cells were pipetted incubated at 80°C for 30min using a thermocycler. The solution of extracted DNA was used as the DNA template for the NGS analysis.

### rhAmpSeq analysis

For each target (EMX1, VEGFA and AAVS1), a PCR primers panel was designed using IDT’s rhAmpSeq designated tool (**Supplementary file 1**)^40^. Library preparation and NGS were conducted according to IDT’s protocol^40^. CRISPResso2^65^ was used to analyze NGS data and assess the on and off-target activity of the Cas variants for each gRNA (using the CRISPRessoPooled utility). Primer panels for EMX1 and VEGFA site 3 were designed according to GUIDE-seq and TTISS data as previously reported^20^. The AAVS1 panel was designed based on GUIDE-seq data from IDT’s rhAmpSeq protocol^40^. The reported off-targets data used in this study were based on experiments conducted under the same conditions (i.e., cell line and gRNA sequence). Each genomic DNA sample was used to assess both on and off target activity.

### GFP disruption assay

EGFP-PEST stable cells were co-transfected with one of the sgRNA plasmids and one of the Cas plasmids, together with a reporter plasmid. Five days post-transfection cells were trypsinized, incubated 5 min at 37 °C, and neutralized by 150ul FBS-enriched FACS buffer (PBS; 5% FBS; 25mM HEPES Biological industries (Israel) 03-025; 5mM EDTA, Biological industries (Israel) 01-862). The suspended cells were filtered using a mesh-capped plate (Merck MANMN4010) before flow cytometry analysis. Cells were analyzed by flow cytometry (CytoFLEX S, Beckman Coulter) to assess GFP-disruption rates. Decrease in GFP levels was assessed by measuring the GFP positive cells, based on GFP+ untreated cells, out of mCherry positive cells (**Sup. Fig. 6**). The GFP disruption rate was calculated as following:

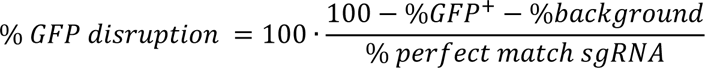

Each plate contained its internal positive and negative controls for normalization.

### Protein extraction and western blot

For the experiments assessing the variants’ protein levels, transfected cells were harvested 72 hours post transfection and treated with RIPA buffer (#89901, Thermo Fisher Scientific, Waltham, MA) freshly supplemented with protease and phosphatase inhibitors (1:100, #78441, Thermo Fisher Scientific), incubated 30 min on ice, and centrifuged at 14,000 RPM for 20 min at 4^◦C^. Protein levels were determined using a bicinchoninic acid assay kit (BCA, #A-4503, Thermo Fisher Scientific). Supernatants were stored at −80°C until used. Thereafter, 30µg of protein was separated by electrophoresis through 7.5% polyacrylamide gels (Mini-PROTEAN, #456-8024, Bio-Rad) and transferred to nitrocellulose membranes (#1704159, Bio-Rad) using Trans-Blot Turbo Transfer System equipment (Bio-Rad). The membranes were blocked using SuperBlock (#37535, Thermo Fisher Scientific) with 0.1% Tween 20 (#P1379-100ML, Thermo Fisher Scientific) and probed overnight at 4°C with the primary antibodies: CAS9 (1:1000, MAB #146977, A9-3A3 Cell-Signaling). The membranes were washed three times with Tris buffered saline (#1706435, Bio Rad, CA, USA) and 0.1% Tris Buffer Saline Tween 20 (TBST) and incubated for 1 h at room temperature with secondary antibodies: goat anti-mouse or goat anti-rabbit IRDye® 800CW/680CW (1:10,000, Licor, Lincoln, NE, USA) for one hour at RT. The membranes were then developed with Odyssey Imager CLx (Licor). As a control for protein loading, blots were probed for mouse anti-Glyceraldehyde 3-phosphate dehydrogenase (GAPDH, 1:1000, MAB #2118, Cell-Signaling) using the same procedures. Data were calculated as the ratio of mean target protein intensity to GAPDH intensity. Densitometric analysis of Western blots was performed using Image Studio Lite software (Licor).

For the p53-pathway activation experiments, cells were transfected with 2ug of plasmid with TransLT1 transfection reagent (Mirus) in antibiotic-free medium, following manufacturer’s protocol. 72 hours post-transfection, cells were harvested with RIPA buffer supplemented with a protease inhibitor cocktail (Sigma-Aldrich #P8340). Proteins were resolved on a 12% SDS-PAGE gel. Bands were detected using chemiluminescence (Millipore #WBLUR0500) on Fusion FX gel-doc (Vilber). Antibodies used: p21 (1:1000, #2947 Cell Signaling Technologies) and GAPDH (1:1000, #5147 Cell Signaling Technologies).

### RNA extraction and qRT-PCR

To assess p53 pathway activation, transfected MCF7 cells were harvested using Bio-TRI® (BioLabs) and RNA was extracted following manufacturer’s protocol. cDNA was amplified using GoScript™ Reverse Transfection System (Promega), using oligoDT primers, following manufacturer’s protocol. qRT-PCR was performed using Sybr Green (Invitrogen), and quantification was performed using ΔCT method. All primers used for detection of p53 pathway targets are available in **Sup. File 1**.

### Immunofluorescence and gH2AX foci quantification

Cells were transfected with 2ug of plasmid with TransLT1 transfection reagent (Mirus) in antibiotic-free medium, following manufacturer’s protocol. 72hrs post-transfection, cells were fixed and stained as previously described^66^. Briefly, cells were fixed with 4% PFA for 15min, then permeabilized with Triton X-100 0.5% for 5min at room temperature. Slides were stained against phospho-gH2AX (1:1000, #05-636 Millipore) for 1.5hrs in a humid chamber. After washing with PBS, cells were incubated with Alexa Fluor 555 anti-mouse antibody (1:1000 Cell Signaling Technologies), then stained with DAPI in PBS (1ug/mL) for 3min.

Images were acquired using cellSense Imaging Software (Olympus) and merged using ImageJ. gH2AX foci were quantified on raw TIFF images using CellProfiler software^67^, with the following parameters: nuclei detection (80-400px size range) using Otsu three class threshold method with middle-intensity class to the background; foci detection (3-20px size range) using Otsu three class threshold method with middle-intensity class to the foreground, and minimal intensity threshold of 0.11. Threshold correction factor and method to distinguish clumped objects were adjusted for each experiment. Detected foci are then attributed to the related detected nuclei.

### Statistical tests

The GraphPad Prism v9.4.1 software was used to produce all charts and analyze data. Statistical tests were performed as described in the figure legends and P values of ≤0.05 are labeled with a single asterisk (*), in contrast to P values of ≤0.01 (**), ≤0.001 (***), or ≤0.0001 (****); where not indicated, P values are non-significant. In all relevant figure panels, values of mean±SEM are reported, and the exact n value is described in each figure legend.

## ACKNOWLEDGEMENTS

The authors would like to thank Michal Shor for her assistance with Figures preparation. This work was supported by the Israel Science Foundation (grant #1010/16 to F.B.), a generous donation from Varda and Boaz Dotan (to F.B.), the European Research Council (ERC Starting Grant # 945674 to U.B.-D.) and by the Israel Science Foundation (grant #1805/21 to U.B.-D.). R.R. was supported by external Ph.D. scholarships from the ‘‘Dan David Prize” and “Teva BioInnovation Fellowship” and travel awards from the TAU Constantiner Institute and the Gertner Institute. J.Z. was supported by fellowships from the Israeli Ministry for Immigrant Absorption, the Pfizer-Wexler Excellence Scholarship, and the Yoran Institute for Human Genome Research, and by travel awards from the TAU Constantiner Institute and Cancer Biology Research Center. The BPK1520 plasmid was a kind gift from Keith Joung (Addgene plasmid #65777; http://n2t.net/addgene:65777;RRID:Addgene_65777).

## AUTHOR CONTRIBUTION

R.R. and O.S. conceptualized and designed the study. R.R. performed the experimental characterization of the enzymes, analyzed the data and wrote the manuscript. O.S. established and performed the computational analyses. J.Z. performed the p53 and *γ*H2AX experiments and analyzed their data. S.H. performed the western blot experiments of Cas9 levels and assisted on the flow cytometry experiments. N.Z. performed the rhAmpSeq data analysis. S.M. assisted in the flow cytometry experiments. J.Z. and N.Y.T. performed RT-PCR in the p53 experiments. R.R., O.S. and J.Z. drafted the manuscript and revised it together with U.B.-D., F.B. and D.O. U.B.D., F.B. and D.O. supervised the study.

---

**Supplementary Figure 1:**
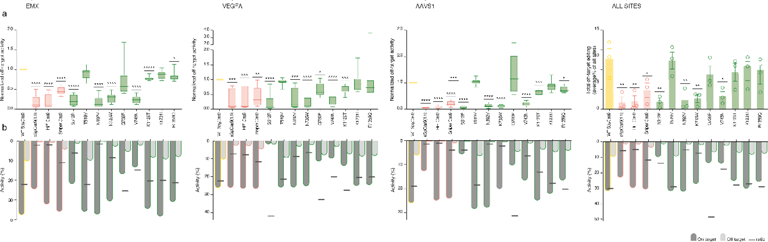
On- and off-target activity of the candidate variants in three genomic loci. **a,** Mean off-target activity of all tested variants for three gRNAs; EMX1, VEGFA and AAVS1. For each gRNA:enzyme N=3. Number of off-target sites for each gRNA: N=8 (EMX1), N=11 (VEGFA) and N=8 (AAVS1). One-way ANOVA, Geisser-Greenhouse correction, Dunnett’s multiple comparisons tests. *p<0.05, **p<0.005, ***p<0.0005, ****p<0.00005. **b,** Representation of the off/on target ratio calculated as 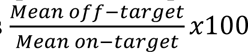.

**Supplementary Figure 2:**
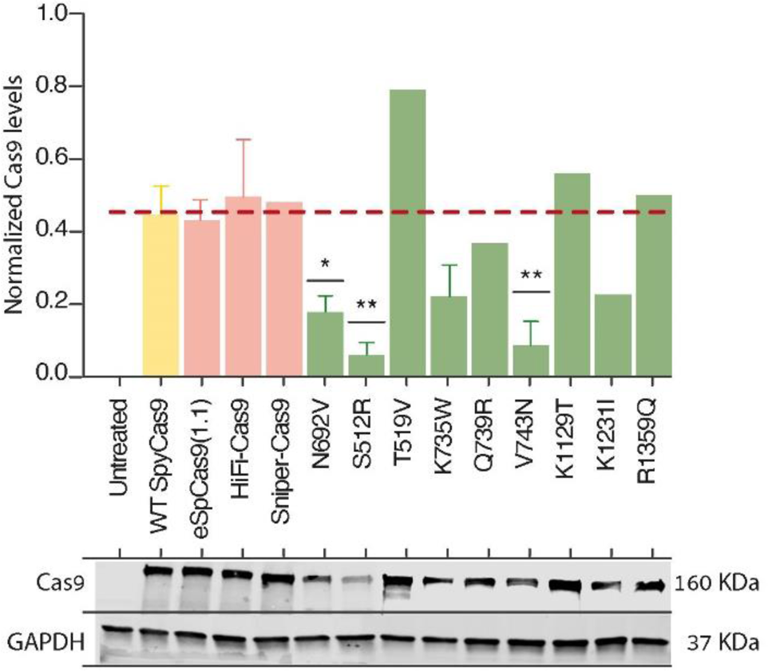
Protein expression levels of HF Cas9 variants. Above are the protein levels of each Cas9 variant normalized to GAPDH. For WT SpyCas9 n=4, eSpCas9(1.1), HiFi-Cas9, S512R and N692V n=3, Sniper-Cas9, T519V, Q739R, K1129T, K1231I and R1359Q n=1. One-way ANOVA, Multiple comparisons to WT SpyCas9. Below are representative images of the Cas9 and GAPDH bands.

**Supplementary Figure 3:**
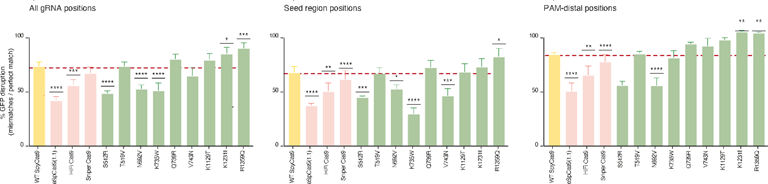
Average GFP disruption of HF variants. Mean values of Fig. 5 heatmap, summarizing the position-specific activity into three analyzes: all positions (1-19), Seed region positions (1-11) and PAM-distal positions (12-19). For each position per variant N=3. One-way ANOVA, Dunnett’s multiple comparisons to WT SpyCas9. *p<0.05, **p<0.005, ***p<0.0005, ****p<0.00005.

**Supplementary Figure 4:**
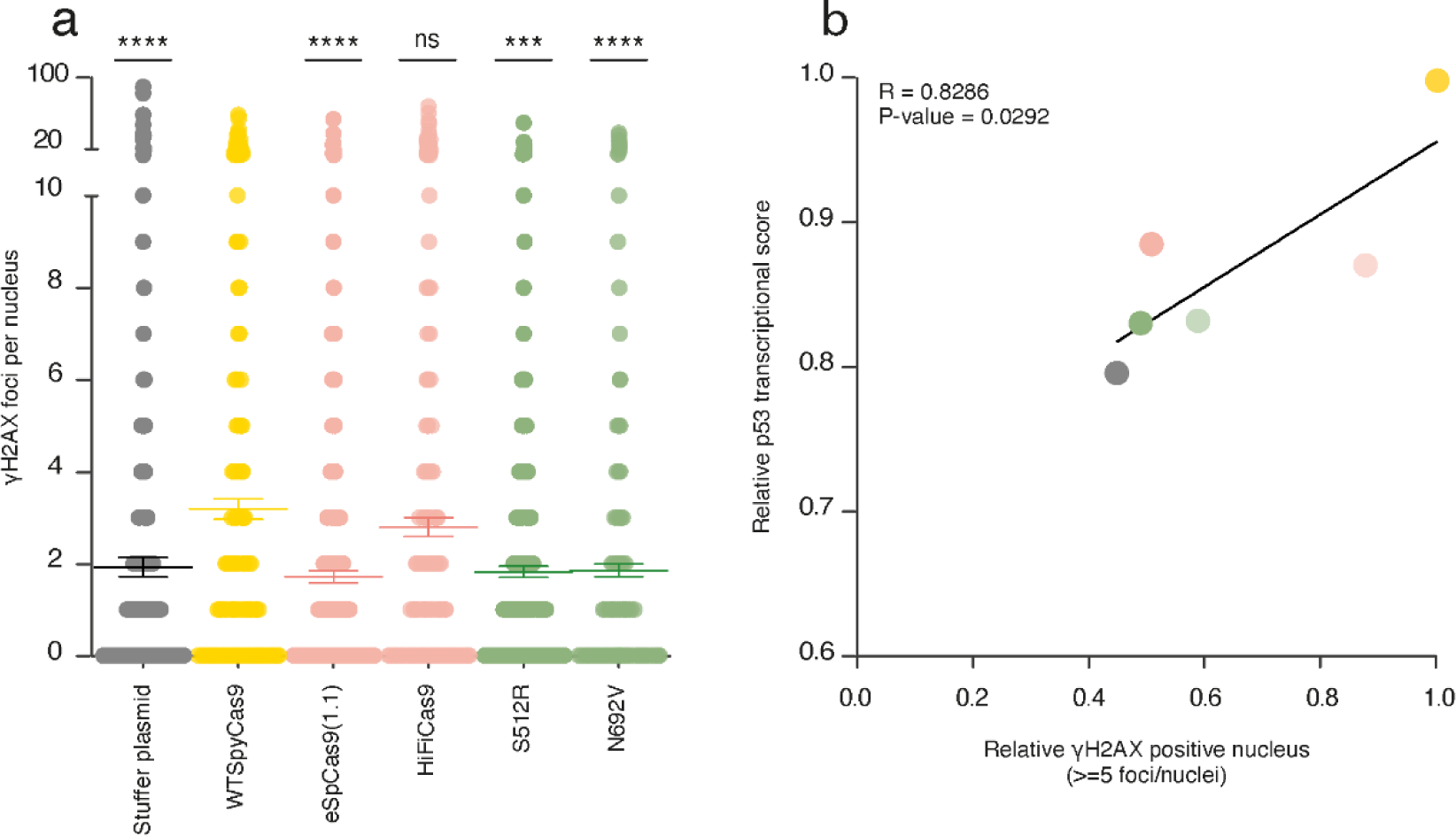
*γ*H2AX foci per nucleus correlate with the relative p53 transcriptional score. **a**, Individual *γ*H2AX foci per nucleus following WT-SpyCas9 or HF variants in MCF7, 72hrs post transfection (raw data from Figure 6). HF variants carried comparable foci per nuclei to control conditions. n=5 independent experiments, about 150 nuclei per condition per experiments; ***p<0.001, ****p<0.0001; Kruskal-Wallis test. **b**, Correlation between p53 transcriptional score (as determined in Figure 6b) and relative *γ*H2AX positive nuclei (as determined in Figure 6f), showing that HF variants are inducing less non-specific DNA damage than WT-SpyCas9. *p=0.0292, rho=0.8286, one-tailed Spearman’s correlation.

**Supplementary Figure 5:**
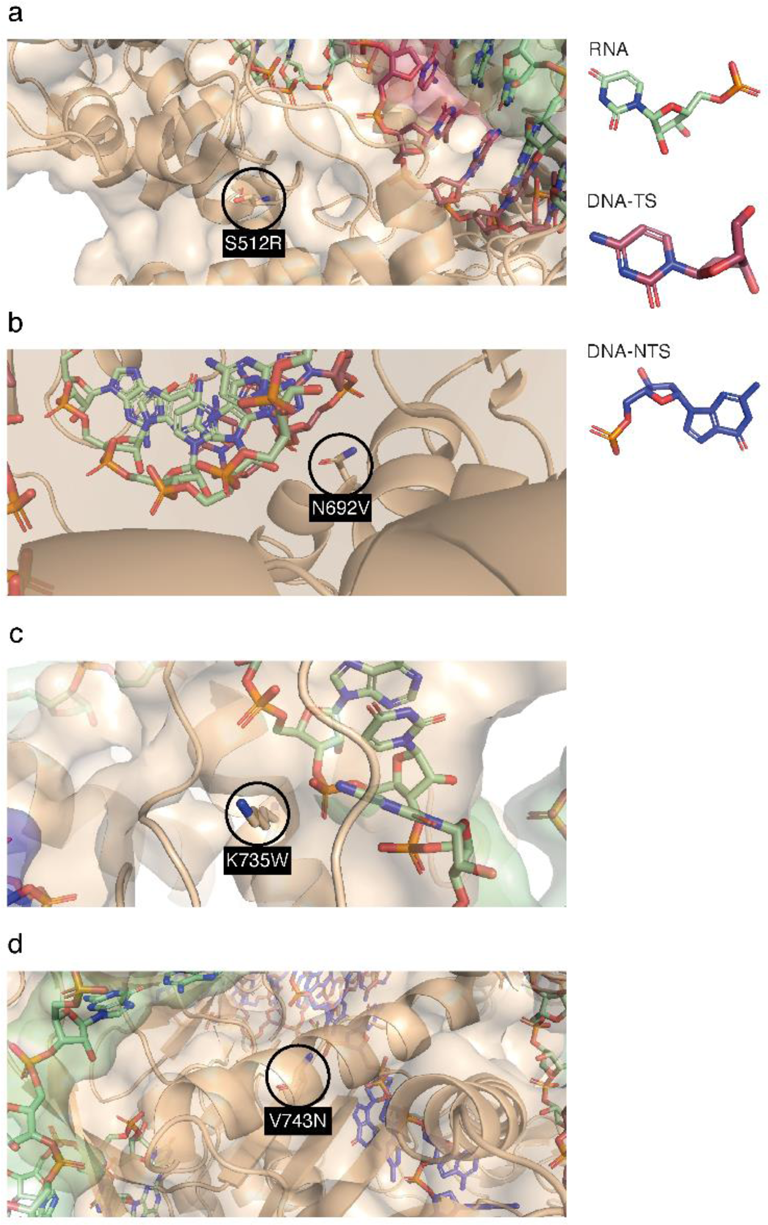
Structural description of the mutated residues of the four HF SpyCas9 variants. RNA nucleotides are shown in green, target strand (TS) DNA in red and non-target strand (NTS) DNA in blue. The protein is represented as cartoon, while the mutated residues (**a**, S512R, **b**, N692V, **c**, K735W and **d**, V743N) are represented as sticks to visualize their distance from nucleic acid in the structure.

**Supplementary Figure 6:**
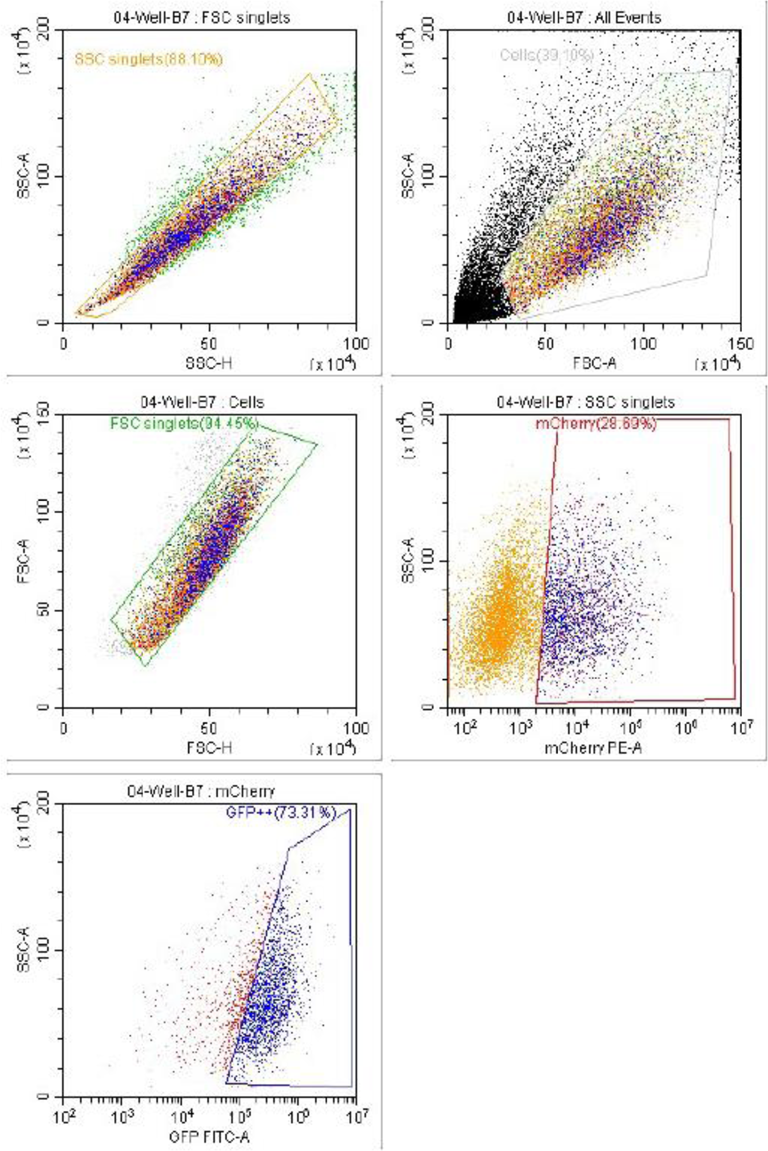
Flow cytometry gating strategy for Fig. 5.

